# Nuclear microenvironments modulate transcription from low-affinity enhancers

**DOI:** 10.1101/128280

**Authors:** Justin Crocker, Albert Tsai, Anand K. Muthusamy, Luke D. Lavish, Robert H. Singer, David L. Stern

## Abstract

Transcription factors regulate gene expression by binding to DNA for short durations and by often binding to low-affinity DNA sequences. It is not clear how such temporally brief, low-affinity interactions can drive efficient transcription. Here we report that the transcription factor Ultrabithorax (Ubx) functionally utilizes low-affinity binding sites in the *Drosophila melanogaster shavenbaby* (*svb*) locus in nuclear microenvironments of relatively high Ubx concentration. By manipulating the affinity of *svb* enhancers, we revealed an inverse relationship between enhancer affinity and Ubx concentration required for transcriptional activation. A Ubx cofactor, Homothorax (Hth), was enriched together with Ubx near enhancers that require Hth, even though Ubx and Hth did not co-localize throughout the nucleus. These results suggest that low affinity sites overcome their kinetic inefficiency by utilizing microenvironments with high concentrations of transcription factors and cofactors. Mechanisms that generate these microenvironments are likely to be a general feature of eukaryotic transcriptional regulation.

## Introduction

Genomic regions near coding genes, called enhancers, direct specific patterns of gene expression^1–3^. Enhancers contain short DNA sequences that bind sequence-specific activating and repressive transcription factor proteins; the integration of these positive and negative signals directs gene expression^4^. Protein-DNA binding is often an ephemeral event; studies in mammalian cells demonstrate that transcription factors disassociate within seconds of binding to DNA^5–10^. Furthermore, recent studies in animals ranging from fruit flies to mammals have revealed that low affinity DNA binding sites are critical to allow related transcription factors to distinguish between binding sites with similar DNA sequences^11–23^. Increasing the affinity of binding sites, to more stably recruit transcription factors, activates promiscuous gene expression^12,24^, which may lead to developmental defects. It is unclear how brief protein-DNA contacts can mediate transcription from enhancers containing low affinity binding sites.

One possible mechanism that could mitigate low binding affinity is an increase in the local concentrations of transcription factors. At the scale of a single enhancer a few hundred base pairs long, multiple low-affinity binding sites for the same transcriptions factor in close proximity could increase the frequency of binding events by locally trapping the protein compared to an isolated binding site. Furthermore, interactions of transcription factors and cofactors with multiple localized enhancers could generate “microenvironments”^2^ of high transcription factor concentrations.

We have explored this problem using the *shavenbaby* (*svb*) locus, which contains multiple enhancers that drive specific patterns of *svb* gene expression that are required for proper development of *Drosophila* embryos. Each of three characterized enhancers contains clusters of low-affinity binding sites for the Hox gene Ultrabithorax (Ubx). These enhancers also require a Ubx cofactor Homothorax (Hth) to function^11^. We have exploited robust transgenic tools in *Drosophila*, new fluorescent dyes, and new approaches to prepare embryos for microscopy to systematically perturb these *svb* enhancers and directly image the results at a sub-nuclear level. We find that microenvironments of high Ubx and Hth concentrations mediate transcription from low-affinity enhancers.

## Results

### Ubx is present in microenvironments of varying local concentrations

We first examined whether nuclei in *Drosophila* embryos possess Ubx microenvironments by performing immunofluorescence (IF) staining in fixed embryos and super-resolution confocal imaging using Airyscan (Carl Zeiss Microscopy, Jena, Germany). We found that Ubx protein was not distributed uniformly, but rather exhibited regions of high and low Ubx intensities (Figure 1a, b). To observe Ubx distribution at higher resolution, we expanded the size of the embryos^25^ by approximately four-fold in each dimension (Figure 1c, materials and methods). Nuclei of expanded embryos revealed distinct regions of high Ubx intensity separated by regions of low Ubx intensity. We observed, on average, 184.9 ± 24.6 (*n* = 12, 3 embryos) clusters per nucleus that were stronger than one-quarter of the maximum Ubx intensity (Figure 1d, e, & S1).

**Figure 1:**
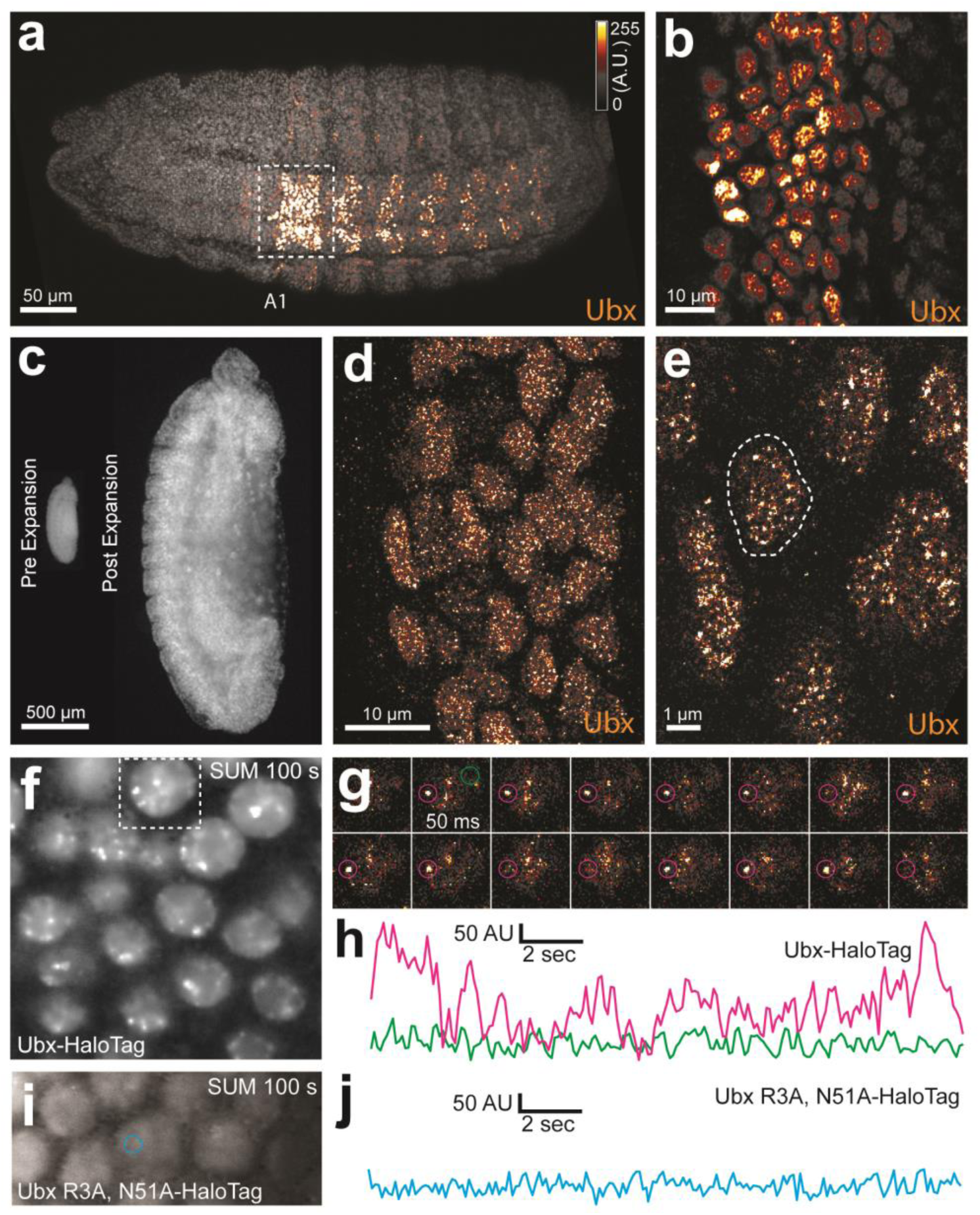
Ubx is present in microenvironments with varying local concentrations. (**a**, **b**) Stage 15 embryos stained for Ubx protein with a bounding box indicating a ventral region of abdominal segment one (A1). (**b**) Higher magnification, Airyscan image of the region indicated in panel (**a**). (**c**) Stage 15 embryo pre- and post-expansion. (**d**, **e**) Expanded stage 15 embryos stained for Ubx protein. The dashed line encircles a single nucleus (**e**). (**f**, **i**) Projections of summed pixel intensity over 100 seconds from videos of *nos*::GAL4, UAS::HaloTag-Ubx for either a wild-type Ubx (**f**) or a binding deficient Ubx (**i**), imaged with JF_635_ Dye. (**g**) Sixteen individual, 50 millisecond video frames of the nucleus surrounded by a dashed box in panel (**f**). (**h**, **j**) Temporal traces of the signal intensity of the regions noted in panel (**g**) or (**i**). The color of each trace corresponds to the colors of the circles in panels (**g**) and (**i**). AU indicates Arbitrary Units of fluorescence intensity.

One explanation for the observed distribution of Ubx is that transcription factors localize generally to accessible regions of the nucleus that have high levels of transcriptional activity. This mechanism, if shared by transcription factors in general, should yield Ubx distributions that mostly overlap with that of other transcription factors. Engrailed (En), a transcription factor unrelated to Ubx, displayed non-uniform sub-nuclear concentrations, but its distribution only partially overlapped that of Ubx (Figure S2a-c, white regions indicate overlap). We similarly observed only partial overlap between Ubx and Even-skipped (Eve) (Figure S2d-f). Abdominal-A (AdbA), a paralog of Ubx that is expressed mainly in separate cells from Ubx and that has similar DNA binding specificity as Ubx, was excluded from Ubx regions in the few nuclei where both were expressed (Figure S3g-i). These results indicate that the distributions of these transcription factors do not result from a shared mechanism that limits the distribution of all transcription factors to the same sub-nuclear space.

We also examined whether Ubx simply occupies regions containing actively transcribed DNA. Both active RNA Polymerase II (Pol II) and the methylated histone H3K4me3, which marks actively transcribed DNA, only partially overlapped with Ubx (Figure S3a-f). In contrast, the histone mark H3K27me3, which marks regions of repressed chromatin, displayed almost no overlap with the distribution of Ubx (Figure S3g-i). Thus, Ubx is not simply restricted to regions inside the nucleus that are available to other transcription factors or to regions of high transcriptional activity.

### Ubx repeatedly binds to specific regions in nuclei of live embryos

We tested whether the heterogeneous protein distributions we observed were an artifact of the fixation protocol^26^ by examining the spatiotemporal dynamics of single Ubx molecules in live *Drosophila* embryos (Figure S4a, material and methods). Single-molecule imaging has been performed previously in cell-lines because live-imaging studies in embryos requires overcoming several new challenges, including imaging at lower signal-to-noise ratios than in cells, compensating for rapid changes during embryonic development, and determining how to deliver fluorescent dyes. We overcame these challenges by generating a HaloTag-Ubx transgene that allowed precise control of fusion protein levels (Figure S4a) and coupling HaloTag-Ubx *in vivo* to new, strongly fluorescent dyes^27^. Over-expression of the *HaloTag-Ubx* transgene recapitulated known embryonic developmental defects, indicating that the HaloTag-Ubx protein retained its function to activate transcription (Figure S4d & e). We then expressed HaloTag-Ubx with a *nanos* promoter, which resulted in HaloTag-Ubx expression in all cells at early developmental stages, and injected the HaloTag ligand of Janelia Fluor 635 (JF_635_)^28^ into these live embryos. JF_635_ is minimally fluorescent in solution but its fluorescence increases by over 100-fold when bound to a HaloTag protein, allowing the detection of labeled Ubx molecules against a background of dim freely diffusing dyes. The fluorescence intensity of labeled Ubx scaled with distance from the site of dye injection (Figure S4b & c), consistent with dye diffusion from the site of injection.

To measure the density of HaloTag-Ubx in nuclei of live embryos, we calculated the average intensity over 100 s (2000 frames). We observed regions of Ubx signal (3-10x background) similar to the high-intensity clusters observed in fixed embryos (Figure 1f). We examined the dynamics of HaloTag-Ubx in nuclei by plotting fluorescence intensity over time (Figure 1g & h and Figure S6). This revealed binding of HaloTag-Ubx in specific nuclear domains with residence times on the order of seconds. These timescales are consistent with transcription factor-DNA binding dynamics measured in live-cell imaging experiments using mammalian cell lines^5,7–10,29^. These repeated binding events apparently produced the high intensities observed in the time-averaged projections. These live-imaging results indicate that Ubx concentrates and remains within specific nuclear regions.

To determine whether regions of high Ubx concentration depended on DNA binding, we performed the same experiments with a version of the *HaloTag*-*Ubx* transgene in which Arg3 and Asn51 of the homeodomain were mutated to Ala (R3A and N51A), which abrogates DNA binding^30^. The DNA-binding deficient Ubx did not display any spatial heterogeneity nor fluctuations in intensity consistent with transcription-factor DNA binding events (Figure 1i & j), suggesting that binding of Ubx to DNA is required to generate restricted nuclear distributions of Ubx.

### Transcriptionally active *svb* loci and enhancers correlate with regions of high Ubx concentration

The heterogeneous distributions of Ubx we observed are consistent with the hypothesis of nuclear “microenvironments”^2^, whereby high local concentrations of transcription factors may drive transcription. Therefore, we examined whether these regions of high Ubx concentration co-localized with sites of active transcription. The *svb* gene is directly regulated by Ubx protein through binding of Ubx to low-affinity sites in multiple *svb* enhancers^11^. We marked sites of active *svb* transcription by fluorescence *in situ* hybridization (FISH) and compared the localization of actively transcribed *svb* loci to Ubx protein concentration (Figure 2a & b). We observed high local Ubx concentrations surrounding active *svb* transcription sites (Figure 2c-f). To quantify Ubx distributions around these sites, we calculated the average Ubx intensity as a function of distance *r* from the point of maximum FISH intensity for each *svb* transcription site (Figure 2g-i and materials and methods). Ubx intensity was normalized to 1 at *r* = 0 (maximum FISH intensity) and averaged across all sites measured. To adjust for background fluorescence, we located the minimum intensity in the averaged Ubx distribution (r = 2-4 μm) and subtracted that value from the distribution. The first μm of the radially averaged 3D distribution is shown, with the shaded area representing the variance (Figure 2j). Within the first μm, *svb* transcription sites showed a relative enrichment of Ubx and were located on average at a local maximum of Ubx concentration. The normalized Ubx intensity after background subtraction at the site of *svb* transcription was 0.60 ± 0.17 (*n* = 59, 4 embryos, uncertainty is the variance of the background) and decreased approximately 250 nm away from the site. Thus, active *svb* transcription sites colocalized with areas of high Ubx concentration spanning approximately a few hundred nanometers.

**Figure 2:**
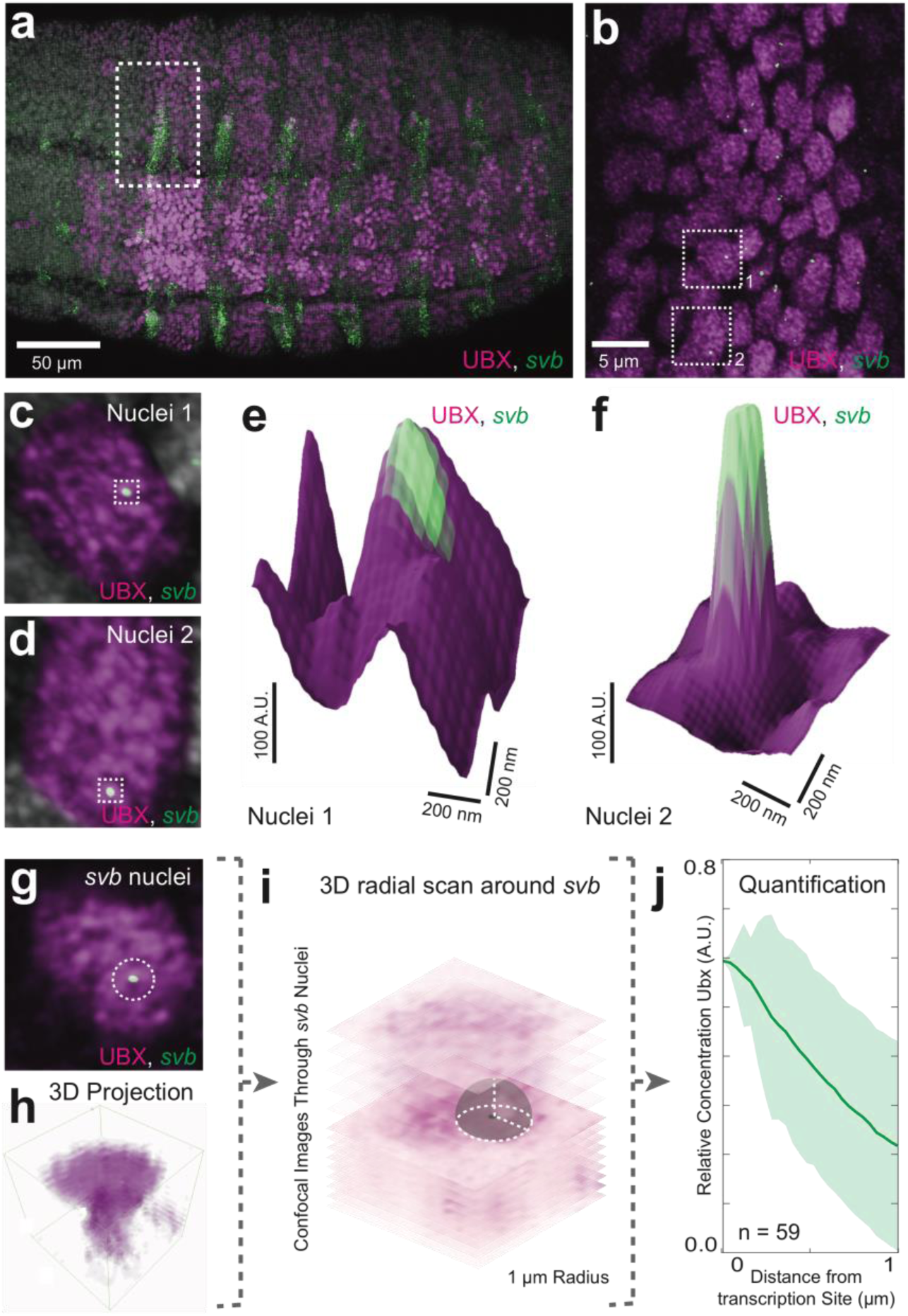
Transcriptionally active *svb* loci reside in regions of high Ubx concentration. (**a**-**d, g**) Embryos co-stained for both Ubx protein and *svb* intronic mRNA. Bright spots of *svb* intronic mRNA mark actively transcribed *svb* loci. (**b**) Higher magnification, Airyscan image of the region noted in panel (**a**), revealing sites of *svb* transcription (green). (**c, d**) Higher magnification, Airyscan images of the nuclei noted in panel (**b**). (**e, f**) 3D surface plots of the images in panels (**c**) and (**d**), centered on the sites of *svb* transcription (green), where height represents Ubx intensity. (h) 3D view of the confocal stack from the nuclei in panel (**g**). (**i**) Schematic outlining the method of Ubx quantification surrounding *svb* transcriptional sites. A 3D radial distribution of the average Ubx intensity on the surface of a sphere centered at the site of *svb* transcription, was calculated. The gray sphere and white outlines is an example of the sphere with a radius *r* = 1 μm. (**j**) Quantification of the average relative concentration of Ubx and the distance from *svb* transcription sites (n = 59, see supplemental methods “settings for extracting radially averaged distributions” for how relative concentration is computed). The shaded region indicates the variance. A.U. indicates Arbitrary Units of fluorescence intensity.

If Ubx protein co-localizes with actively transcribed *svb* loci because Ubx drives *svb* expression, then we would expect that transcription at a locus not regulated by Ubx should not co-localize with high Ubx concentrations. Indeed, we observed that active transcription sites driven by a synthetic enhancer containing binding sites for a TALEA transcription factor^4,31,32^ did not show Ubx enrichment on average despite wide fluctuations in Ubx levels, with a relative enrichment of Ubx at TALEA driven enhancers of 0.02 ± 0.63 (Figure 3a-c, *n* = 29, 3 embryos).

**Figure 3:**
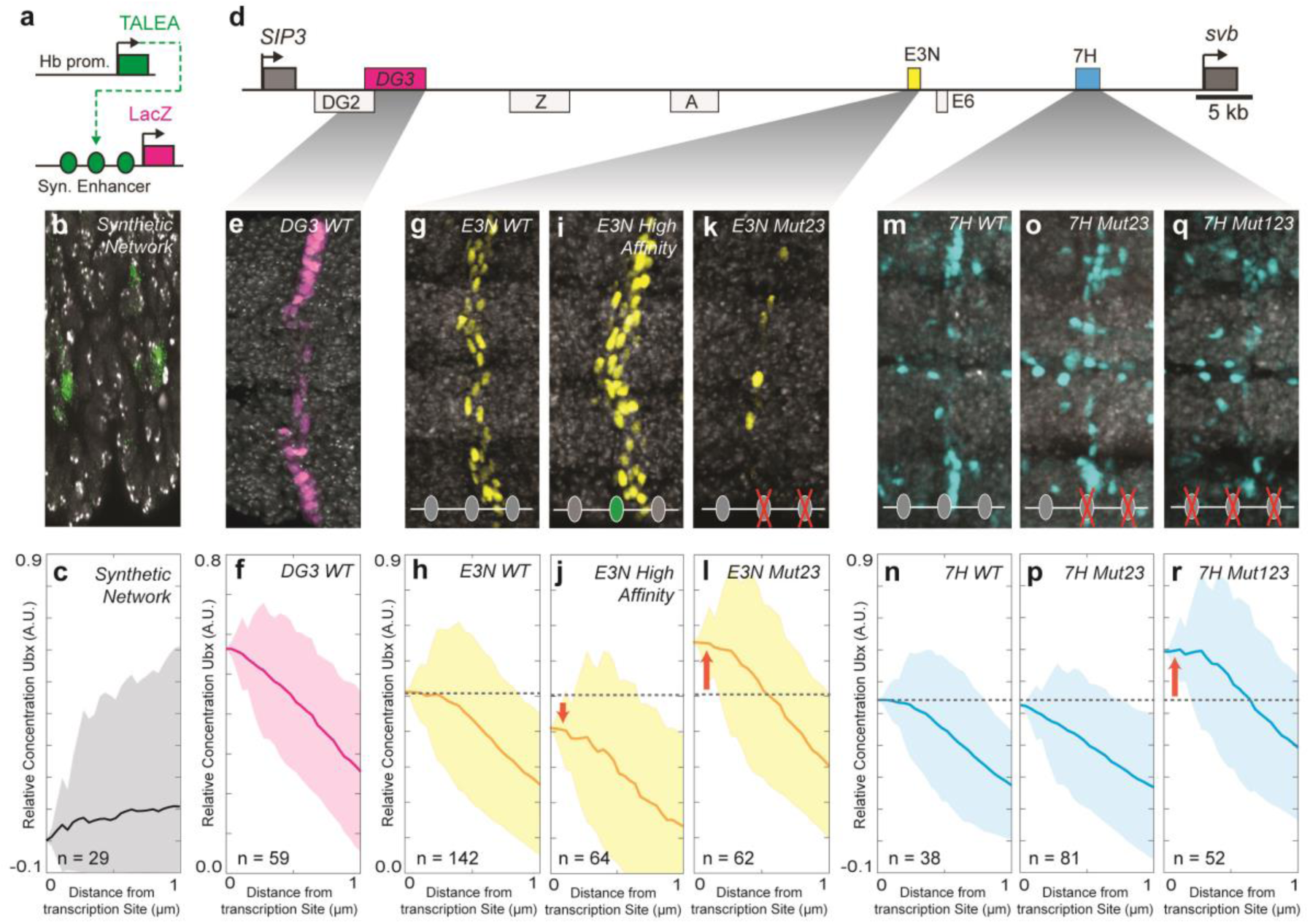
Manipulation of Ubx binding site number and affinity alters the level of Ubx enrichment around *svb* enhancers. (**a**) Schematic of the synthetic TALEA transcription network driven by the Hunchback (Hb) promoter, indicating TALEA binding sites with green circles. (**b**) Early stage 15 embryos carrying the TALEA synthetic network stained with an antibody against β-Galactosidase. (**c**) Quantification of the relative concentration of Ubx based on the distance from synthetic network transcription sites. (**d**) Schematic of the *shavenbaby* locus, indicating embryonic *cis*-regulatory enhancers in boxes. The ventral embryonic enhancers *DG3*, *E3N* and *7H* are highlighted in magenta, yellow and blue boxes, respectively. (**e, g, i, k, m, o, q**) Early stage 15 embryos carrying the reporter constructs *DG3-lacZ* (**e**), *E3N-lacZ* (**g, i, k**), or *7H-lacZ* (**m, o, q**) stained with an antibody against β-Galactosidase, with Ubx-Exd sites altered as indicated. (**f, h, j, l, n, p, r**) Quantification of the relative concentration of Ubx versus the distance from *svb* transcription sites. The shaded regions in panels (**c, f, h, j, l, n, p, r**) indicate the variance. A.U. indicates Arbitrary Units of fluorescence intensity.

### Manipulation of binding site number and affinity changes the level of Ubx enrichment around *svb* enhancers

The experiments described so far showed that the actively transcribed native *svb* locus co-localizes with local concentration maxima of Ubx in the nucleus. We wondered whether binding site affinity modulates the location of transcription relative to Ubx microenvironments. To address this question we examined transcription driven by the individual *svb* enhancers *DG3*, *E3N*, and *7H*, each of which contains a cluster of low-affinity Ubx binding sites and can independently drive transcription of a reporter gene when moved from their native location^11^. Transcription sites driven by these relocated enhancers also colocalized with regions of high Ubx concentration (Figure 3d). The relative Ubx enrichment for each of the three enhancers was 0.56 ± 0.16 for *DG3* (*n* = 61, 3 embryos), 0.51 ± 0.19 for *E3N* (*n* = 142, 11 embryos), and 0.68 ± 0.10 for *7H* wildtype (*n* = 38, 3 embryos) (Figure 3e-h, m, n). These results indicate that low-affinity enhancers actively transcribed far from the native *svb* locus also co-localize with microenvironments of high Ubx concentrations.

Increasing the binding affinity of a site should increase its sensitivity to lower Ubx concentrations. We found previously that replacing a single low-affinity Ubx site with one of a higher affinity led to higher levels of expression and sometimes drove promiscuous transcription^11^, suggesting that more stable Ubx-DNA interactions allowed higher transcriptional activation. Consistent with these previous results, we observed that increasing the affinity of a single low-affinity binding site in the *E3N* enhancer decreased enrichment in Ubx microenvironments to 0.44 ± 0.27 (Figure 3i & j, *E3N High Affinity*, *n* = 36, 3 embryos).

In contrast, we reported previously that deletion of low affinity binding sites reduced transcription^11^. Removing some Ubx binding sites should lower the effective affinity of the entire enhancer and we hypothesized that this may result in transcription only when genes are associated with areas of higher Ubx concentrations. Consistent with this model, when we deleted two low-affinity sites in *E3N*, active transcription was observed in regions of increased Ubx enrichment (0.65 ± 0.18, Figure 3k & l, *E3N* Mut23, *n* = 62, 5 embryos). Deletion of two low-affinity Ubx sites for the *7H* enhancer did not alter Ubx enrichment around transcription sites (0.63 ± 0.37, Figure 3o & p, *7H* Mut23 *n* = 81, 6 embryos). But, deletion of three Ubx binding sites in the *7H* enhancer increased relative Ubx enrichment, consistent with the pattern we observed for the *E3N* enhancer (0.91 ± 0.27, Figure 3q & r, *7H* Mut123, *n* = 52, 8 embryos).

Across all manipulations, we observed an inverse correlation between affinity and the distribution of Ubx intensities at transcription sites (Figure S7). Thus, the number of Ubx binding sites and their affinities determine the response of *svb* enhancers to local Ubx concentration. Lower affinity enhancers require higher Ubx concentrations to drive transcription. Conversely, higher affinity enhancers can drive transcription at lower local Ubx concentrations.

### The Ubx cofactor Homothorax (Hth) is co-enriched around transcription sites with Ubx

Co-factors can stabilize low-affinity binding interactions through cooperative interactions with transcription factors. A co-factor-dependent enhancer would require sufficient concentrations of both the factor and the co-factor to drive transcription. The homeodomain proteins Extradenticle (Exd)/Pbx and Homothorax (Hth)/MEIS^30,33–35^ interact with Ubx during DNA binding and Ubx and Hth regulate a partially overlapping set of genes^36,37^. *In vitro*, Ubx requires Hth/Exd to bind to the low-affinity sites in *7H* and *E3N*^11^. *In vivo*, Hth deficiency led to the loss of expression for both *7H* and *E3N* (Figure 4a-d). Consistent with this co-requirement for Ubx and Hth, Hth was co-enriched with Ubx around active transcription sites driven by *7H* or *E3N* (Figure 4e-t). The relative enrichment for Ubx and Hth respectively was 0.58 ± 0.14 and 0.41 ± 0.16 for *7H* (*n* = 51, 7 embryos) and 0.66 ± 0.13 and 0.39 ± 0.24 for *E3N* (*n* = 74, 5 embryos). These results suggest that transcription from co-factor-dependent enhancers requires microenvironments that contain high concentrations of both transcription factors and their co-factors.

**Figure 4:**
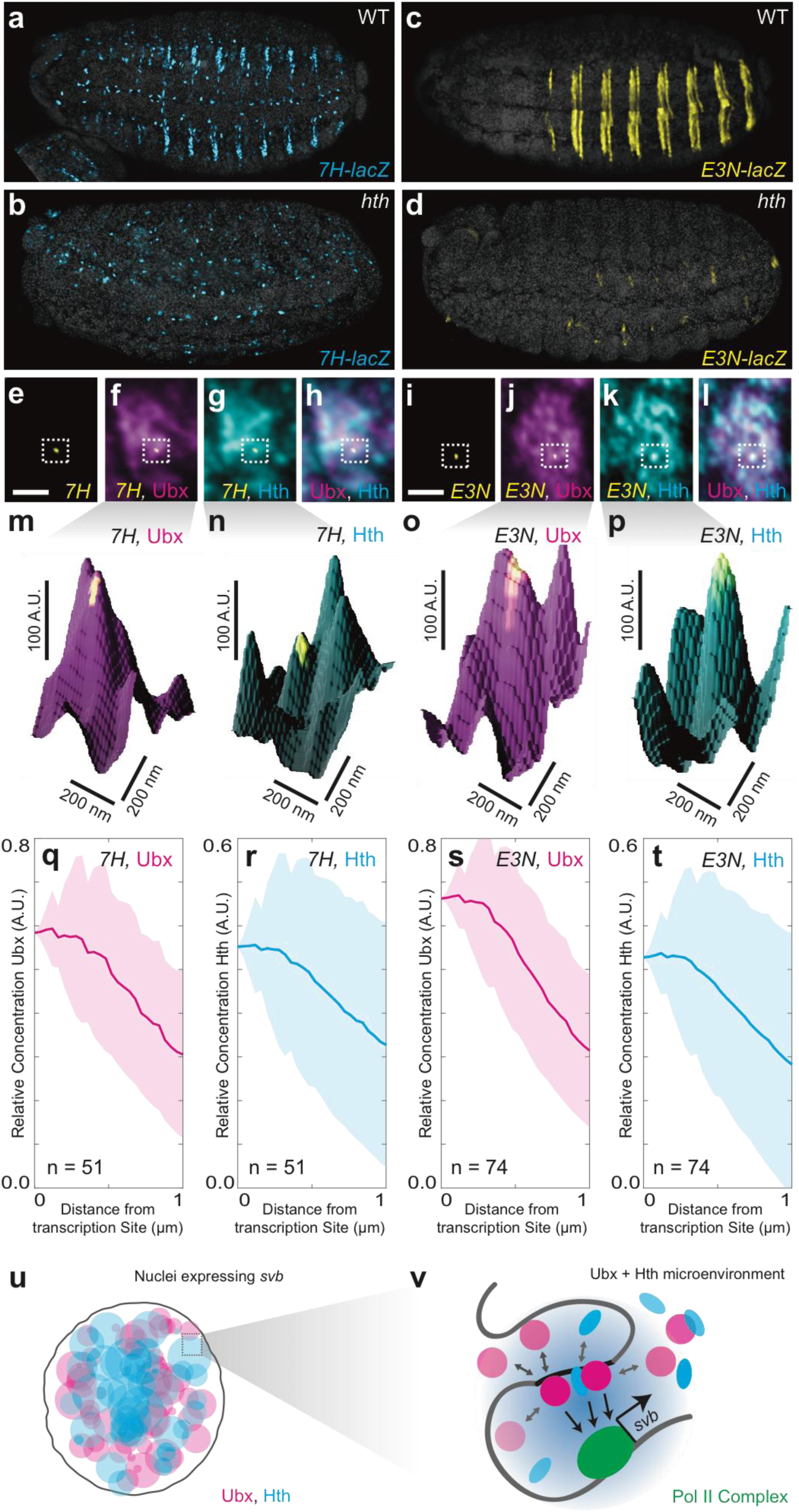
Ubx and its cofactor Hth are co-enriched around transcription sites. (**a-d**) Early stage 15 embryos with *7H-lacz* (**a-b**) or *E3N-lacZ* reporter constructs (**c-d**) stained with an antibody against ß-Galactosidase in either Wild-type (WT) (**a** and **c**) or *hth^P2^* mutant embryos (**b** and **d**). (**e-h**) A nucleus displaying active transcription of the *7H-lacZ* reporter construct denoted by a bounding box (**e-h**) and co-stained for Ubx protein (**f**), Hth protein (**g**), or both Ubx and Hth proteins (**h**). (**i-l**) A nucleus displaying active transcription of the *E3N-lacZ* reporter construct denoted by a bounding box (**i-l**) and co-stained for Ubx protein (**j**), Hth protein (**k**), or both Ubx and Hth proteins (**l**). (**m-p**) 3D surface plots of the images in panels (**f, g, j, k**), centered on the sites of enhancer activity (yellow). The height of the plot is Ubx intensity in panels (**m**) and (**o**) and Hth intensity in panels (**n**) and (**p**). (**q-t**) Quantification of the relative concentration of Ubx (**q, s**) and Hth (**r, t**) versus distance from active enhancer sites. The shaded regions indicate the variance. A.U. indicates Arbitrary Units of fluorescence intensity. (**u, v**) A conceptual model showing nuclei with multiple regions of high local concentrations of Ubx or Hth (**u**) and high local concentrations of both Ubx and Hth that allow rapid ON rates (**v**, grey arrows) and collectively may recruit RNA pol II complexes.

## Discussion

Biological systems often generate locally high concentrations of interacting molecules to increase the efficiency of biochemical reactions^38,39^. This appears to be true also for transcription from low-affinity enhancers. Microenvironments^2^ of high local concentrations of transcription factors and their co-factors may circumvent the instability of low-affinity interactions by promoting more frequent DNA binding and cooperative interactions^13^ (Figure 4u & v). Many mechanisms might create these observed transcription factor microenvironments. First, clustered binding sites for the same transcription factor^18^ could lengthen the dwell time of proteins near enhancers and increase effective local protein concentrations^40–45^. Second, cooperative interactions between transcription factors and co-factors, each of which may bind independently to enhancers, may stabilize transcription factors at low-affinity sites^13^,^46^. Finally, clustering of enhancers could trap transcription factors over longer length scales^47–51^, perhaps generating the 200 to 400 nm microenvironments that we observed. This last model is supported by recent findings that multiple promoters can share the same enhancer in a common local environment^52^.

Transcription factor microenvironments may be a general feature of eukaryotic transcription, as supported by studies showing mouse and human cells exhibiting RNA polymerase II crowding^53^,^54^, transcription factors using local clustering to efficiently find their binding sites^5^,^7^, and chromatin packaging in *Drosophila* cells generating distinct chromatin environments at the kilobase-to-megabase scale^55^. Collectively, these findings are consistent with a phase-separated model of transcriptional regulation^56^, whereby transcription occurs in distinct microenvironments containing the correct combination of proteins. Enhancers acting as DNA scaffolds for protein binding could provide the anchoring interactions that form transcriptional microenvironments. These microenvironments could, in turn, provide a mechanism to allow efficient transcription from low-affinity enhancers.

## Acknowledgements

We thank Richard Mann, Timothée Lionnet, Paul Tillburg and Brian English for experimental design advice and assistance. We thank François Payre for advice on data presentation. We thank all members of the Stern and Singer labs for discussion. Albert Tsai is a Damon Runyon Fellow of the Damon Runyon Cancer Research Foundation (DRG 2220-15). Robert H. Singer is supported by the 4D Nucleome Award U01-EB21236.

## Materials and methods

### Preparing fixed *Drosophila* embryos

*D. melanogaster* strains were maintained under standard laboratory conditions. All enhancer constructs were cloned into the placZattB expression construct with a hsp70 promoter^1^. Transgenic fly lines were made by Rainbow Transgenic Flies Inc. *E3* and *7H* were integrated at the attP2 landing site. *DG3* was integrated at ZH-86Fb.

### Immuno-fluorescence staining of transcription factors and *in situ* hybridization to mRNA

Flies were reared at 25 °C and embryos were fixed and stained according to standard protocols^1^. Primary antibodies were detected using secondary antibodies labeled with Alexa Fluor dyes (1:500, Invitrogen). In situ hybridizations were performed using DIG labeled, antisense RNA-probes against reporter construct RNA or the first intron of *svb*. DIG RNA products were detected with a DIG antibody: Invitrogen, 9H27C19 (1:200) or LacZ: Promega anti-ß-Gal antibody (1:1000).

The following primary antibodies were used at the indicated concentrations:

Ubx: Developmental Studies Hybridoma Bank, FP3.38-C (1:20)
Hth: Santa Cruz Biotechnology (dN-19), sc-26186 (1:50)
Eve: Developmental Studies Hybridoma Bank, 2B8-C (1:20)
AbdA: Santa Cruz Biotechnology (dN-17), sc-27063 (1:50)
En: Santa Cruz Biotechnology (d-300), sc-28640 (1:50)
RNA PolII RPB1: BioLegend, (920304), (1:200)
Histone H3K27me3: Active Motif, 39157 (1:200)
Tri-Methyl-Histone 3K4: Cell-signaling technology C42D8 (1:200)

### Imaging fixed embryos with Airyscan

Fixed *Drosophila* embryos mounted in ProLong Gold mounting media (Molecular Probes, Eugene, Oregon, USA) were imaged on a Zeiss LSM 880 confocal microscope with Airyscan (Carl Zeiss Microscopy, Jena, Germany) using Airyscan in SR mode to obtain images with 1.7-fold higher resolution compared to diffraction-limited confocal imaging^57^ (supplemental methods: imaging setup for Airyscan). Images presented in the figures were processed with ImageJ^58^.

### Expanding fixed embryos

To expand embryos, after fixation and staining, embryos were embedded into poly-acrylate gels and expended according to a previously published protocol^25^ (supplemental methods: handling expansion gels).

### Imaging expanded embryos

Expanded gels containing embryos were imaged in 6-well glass bottom plates (Cellvis, Mountain View, California, USA) using a Zeiss LSM 800 confocal microscope (Carl Zeiss Microscopy, Jena, Germany) using standard settings (supplemental methods: imaging setup for expanded embryos).

### Preparing embryos for live imaging

Embryos were injected following previously established protocols^59^ with the HaloTag ligand of JF_635_. Briefly, embryos were collected for 30 minutes at 25 °C and placed in oxygen permeable Halocarbon 27 oil. The stock dye solution of 1 mM JF_635_ with a HaloTag ligand in DMSO was diluted 1:100 into fly injection buffer and injected into the posterior end of the embryos. The embryos were then aged to stage 8 and imaged in oxygen permeable Halocarbon 27 oil.

### Live imaging of Drosophila embryos

Injected embryos were imaged on a customized inverted Nikon Ti-Eclipse (Nikon Instruments, Tokyo, Japan) with the appropriate settings (supplemental methods: imaging setup for live embryos).

### Radially averaged distributions centered around transcription sites

To obtain the distributions of Ubx and Hth around a transcription site, the processed stacks obtained from the Zeiss LSM 880 confocal microscope were analyzed in Fiji^60^ using native functions and the 3D ImageJ Suite plugin^61^. Radially averaged distributions were computed in Matlab (MathWorks, Natick, Massachusetts, USA) using custom scripts (supplemental methods: settings for extracting radially averaged distributions).

## Supplemental methods

### Imaging setup for Airyscan

All Airyscan images were acquired using a Zeiss Plan-Apochromat 63x/1.4 Oil DIC M27 objective due to its well-characterized point spread function. First an embryo at the appropriate developmental stage (stage 15 for most embryos) and proper orientation was located. The band of mRNA expression in high Ubx regions was then found. Within that band, areas containing transcription sites in nuclei of high Ubx expression were imaged. Images with both Ubx and Hth were acquired in the same manner by locating the proper area using the mRNA and Ubx. When Ubx was imaged together with RNA polymerase II, a histone marker, or other transcription factors Ubx expression was used to locate the region of interest.

The optimal setting suggested by Zeiss for the number of pixels in the x-y direction (40 nm per pixel) and displacement in the z-stack (190 nm) were used for all Airyscan images. The images from different fluorophores were acquired sequentially with the appropriate laser lines (405 nm, 488 nm, 561 nm, or 633 nm) and spectral filters. The laser power and gain were adjusted to maximize the signal to noise ratio within the dynamic range of the Airyscan detector. The acquired stacks were processed with Zen 2.3 SP1 (Carl Zeiss Microscopy GmbH, Jena, Germany) in 3D mode to obtain super-resolved images.

### Handling expansion gels

To allow easier handling of expanded gels, the gels containing embryos were cast into 8-well silicone isolators without adhesives (8 round chambers with a diameter of 9 mm and a thickness of 0.5 mm, Grace Bio-Labs (Bend, Oregon, USA)) and allowed to polymerize. The gels were transferred into a 6-well glass bottom cell culture plate (Cellvis, Mountain View, California, USA) and expanded using ultrapure water containing 500 nM DAPI. Before imaging, the water was removed and the gel encased in 3% low melting temperature agarose (NuSieve GTG Agarose, Lonza Group Ltd, Basel, Switzerland), taking care not to allow the agarose to flow under the gel and float the gel away from the cover glass bottom. Water was then added back into the wells to prevent drying.

### Imaging setup for expanded embryos

A long working-distance water immersion objective, the Zeiss LCI Plan-Neofluar 25x/0.8 Imm Korr DIC M27 from, was selected for index-matching with the gel and the ability to image deeply enough to cover the first few layers of cells from the surface of the embryos. Stage 15 embryos in the correct orientation were located using the DAPI and Ubx staining. Regions of low to high Ubx expression were imaged sequentially using the appropriate laser lines (405 nm, 488 nm, or 561 nm) with the proper spectral filters. The laser power settings and the gain were selected to maximize signal to noise within the dynamic range of the detector. The full field of view of the microscope was imaged with 2048 × 2048 pixels and with a z-step of 1 μm. The final images presented were processed in ImageJ^58^.

### Imaging setup for live embryos

All videos were collected under a Nikon CFI Plan Apo NCG 100X Oil NA 1.41 objective with an Andor iXon 897 EMCCD camera (Andor Technology Ltd., Belfast, UK). Embryos at stage 8 in the corrected orientation were found and imaged. We selected an area in the middle of the embryo with enough dye-labeled Ubx molecules to observe single molecules and we avoided regions close to the injection site to avoid oversaturating the camera (compare with Figure S5a-d where there are too many labeled Ubx). The samples were illuminated with a 633 nm laser to image the JF635 tagged Halo-Ubx molecules with laser power and camera gain set to maximize signal from individual Ubx molecules without oversaturating the EMCCD detector. The 512x512 pixel videos were acquired at an exposure time of 50 ms per frame for 200 s. Images were processed using ImageJ to generate the time-averaged images and the intensity over time traces presented in the figures.

### Settings for extracting radially averaged distributions

To extract radially averaged protein distributions, we used Fiji to identify transcription sites inside nuclei by thresholding at a level that is roughly 50-fold above the background intensity. The center of a transcription site was defined as the pixel of maximum intensity in 3D in the mRNA channel inside a nucleus with high levels of Ubx expression. The radially averaged distribution out to a radius of 4 μm from transcription site for the transcription factor in 3D was computed using the 3D ImageJ Suite. The suit generates the distribution by computing the average intensity on the surface of a sphere with a radius *r* from the center in three dimensions for all the values of *r* ranging from zero to a desired outer limit (4 μm in this case).

Using custom Matlab scripts, the individual distributions from each transcription site were normalized to have the intensity at the center (*r* = 0) equal to 1. The distributions were averaged. To adjust for background Ubx intensity outside of the nucleus, the entire averaged distribution was offset by a constant value to bring the minimum intensity present in the distribution to zero to generate the distribution plots. The shaded area around the line represents the variance. The first μm of the distributions, where contributions from outside of the nucleus were minimal, are shown in the figures. The relative enrichment of Ubx or Hth for each enhancer variant is the intensity at *r* = 0 in the distribution and the cited uncertainty is the variance at the location of zero Ubx or Hth intensity (the site of minimum intensity before offsetting, between 2 to 4 μm from the transcription site).

The initial dataset for *7H* enhancers contained only a part of the deletion series. A subsequent dataset contained all the *7H* deletion mutants. The *7H* mutants present in both sets were compared and the distributions of Ubx intensity between the sets were found to differ by a multiplicative factor. When such factor was computed for each overlapping *7H* mutant present in both datasets, the results were similar, indicating that there was a systematic shift in background noise. This could have resulted from differences in embryo handling during fixation, antibody staining, and other steps in sample preparation. Other characteristics such as the functional form of the distributions between the two sets and the trends between *7H* mutants within each set remained unchanged after correcting for the difference in intensity. The wildtype *7H* data from the first set with a correction factor and the rest of the deletion series uncorrected from the second set were used to minimize the normalization employed.

---

**Figure S1:**
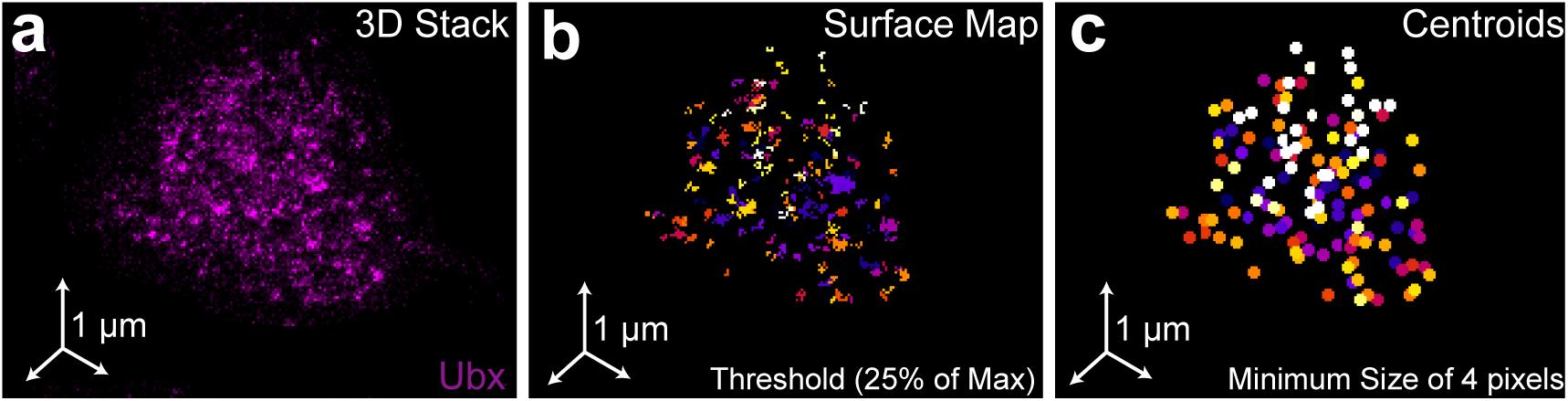
Quantification of Ubx microenvironments in single nuclei. (**a**) 3D projection of a single nucleus stained for Ubx protein. (**b**) Surface plot of contiguous Ubx regions containing a minimum of four pixels with signal intensity greater than twenty-five percent of the maximum intensity. (**c**) Centroids of the Ubx regions found in panel (**b**).

**Figure S2:**
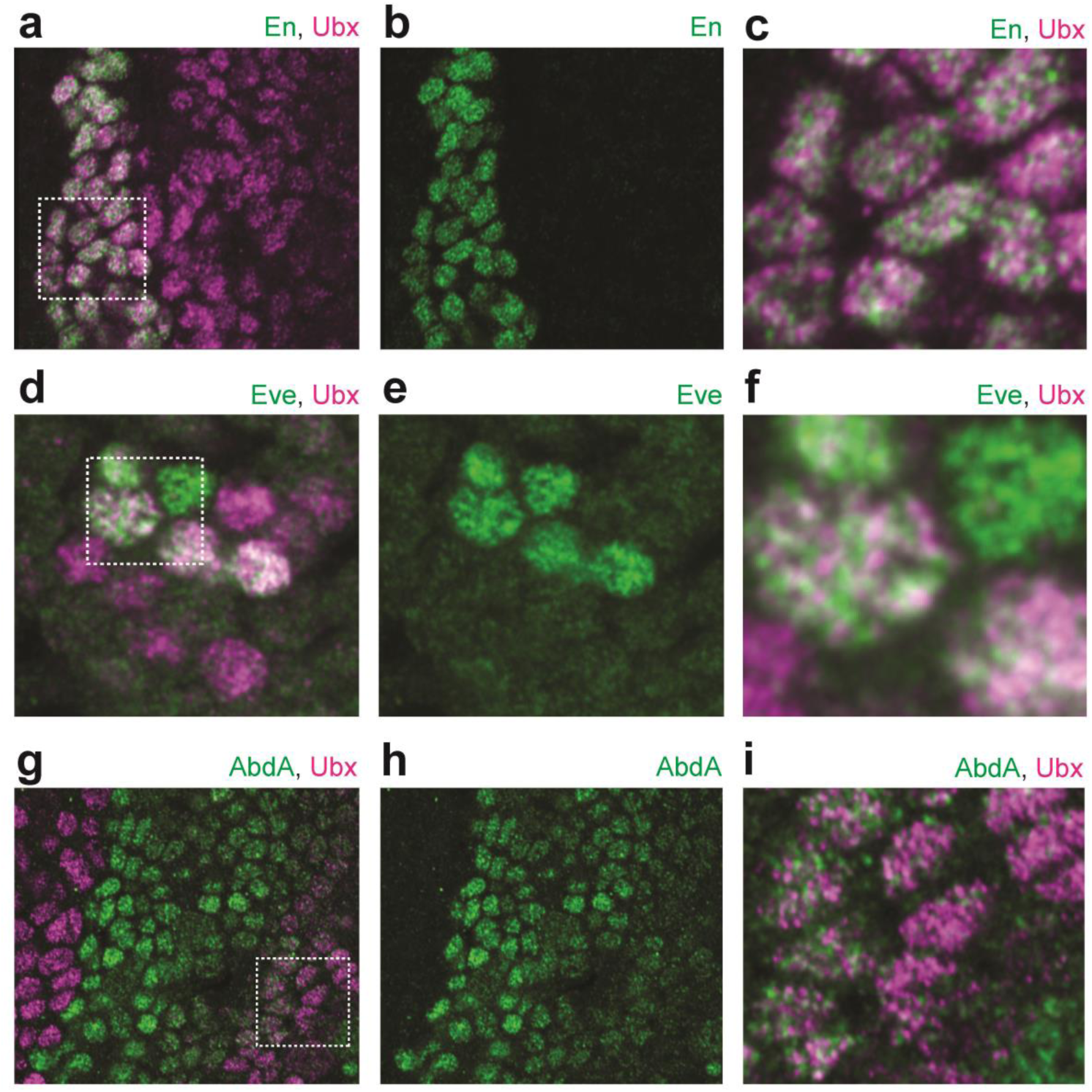
Ubx distribution compared to other transcription factors. (**a-c**) Stage 15 embryo stained with an antibody against Engrailed (En) and Ubx protein (**b** and **c**). (**c**) A higher magnification of the area in the bounding box of panel (**a**). (**d-f**) Stage 15 embryo stained with an antibody against histone Eve and Ubx protein (**e** and **f**). (**f**) A higher magnification of the area in the bounding box of panel (**d**). (**g-h**) Stage 15 embryo stained with an antibody against histone AbdA and Ubx protein (**h** and **i**). (**i**) A higher magnification of the area in the bounding box of panel (**g**).

**Figure S3:**
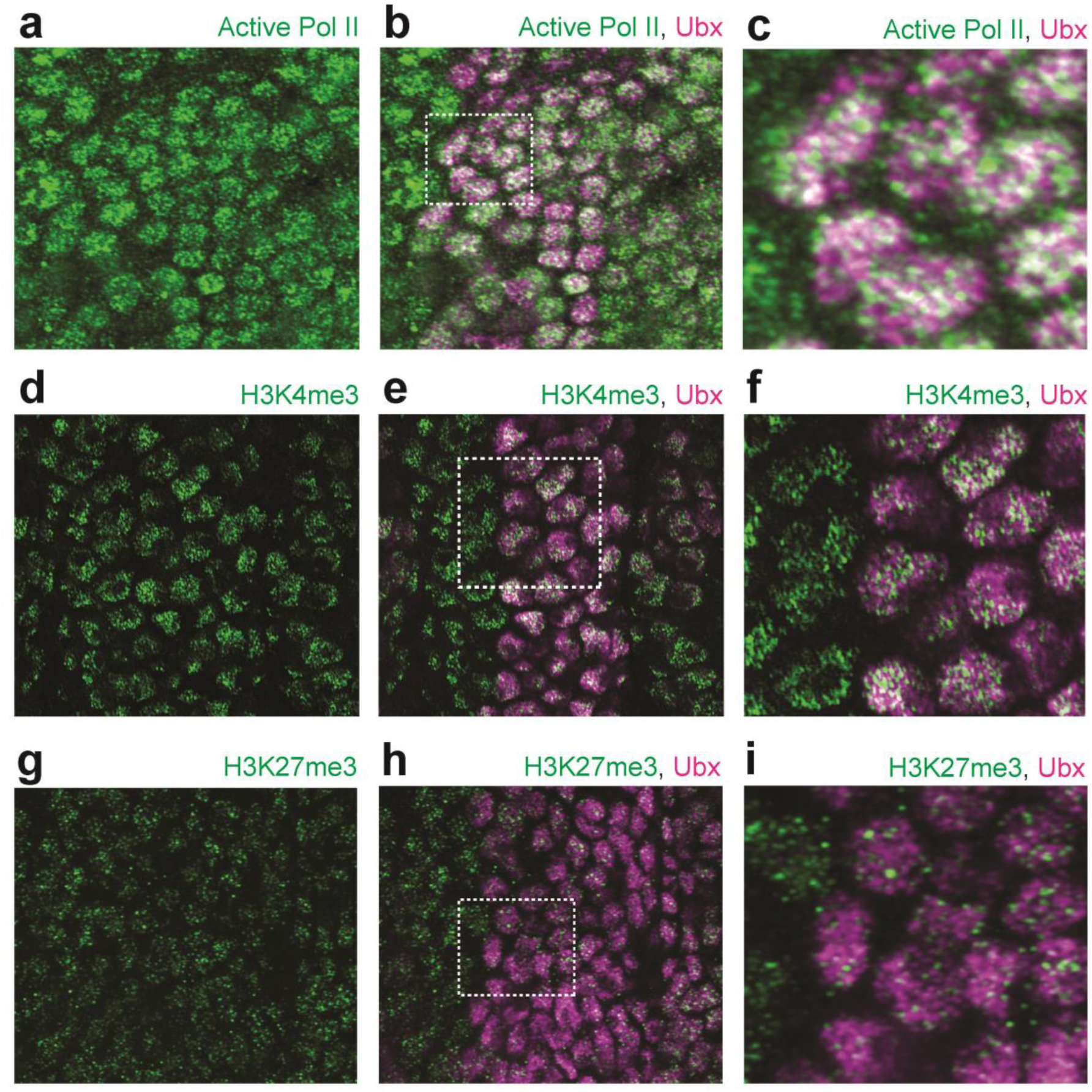
Ubx distribution compared to general markers for transcriptional activity. (**a-c**) Stage 15 embryo stained with an antibody against RNA Polymerase II RPB1 (active Pol II) and Ubx protein (**b** and **c**). (**c**) A higher magnification of the area in the bounding box of panel (**b**). (**d-f**) Stage 15 embryo stained with an antibody against histone H3K4me3 and Ubx protein (**e** and **f**). (**f**) A higher magnification of the area in the bounding box of panel (**e**). (**g-h**) Stage 15 embryo stained with an antibody against histone H3K27me3 and Ubx protein (**h** and **i**). (**i**) A higher magnification of the area in the bounding box of panel (**h**).

**Figure S4:**
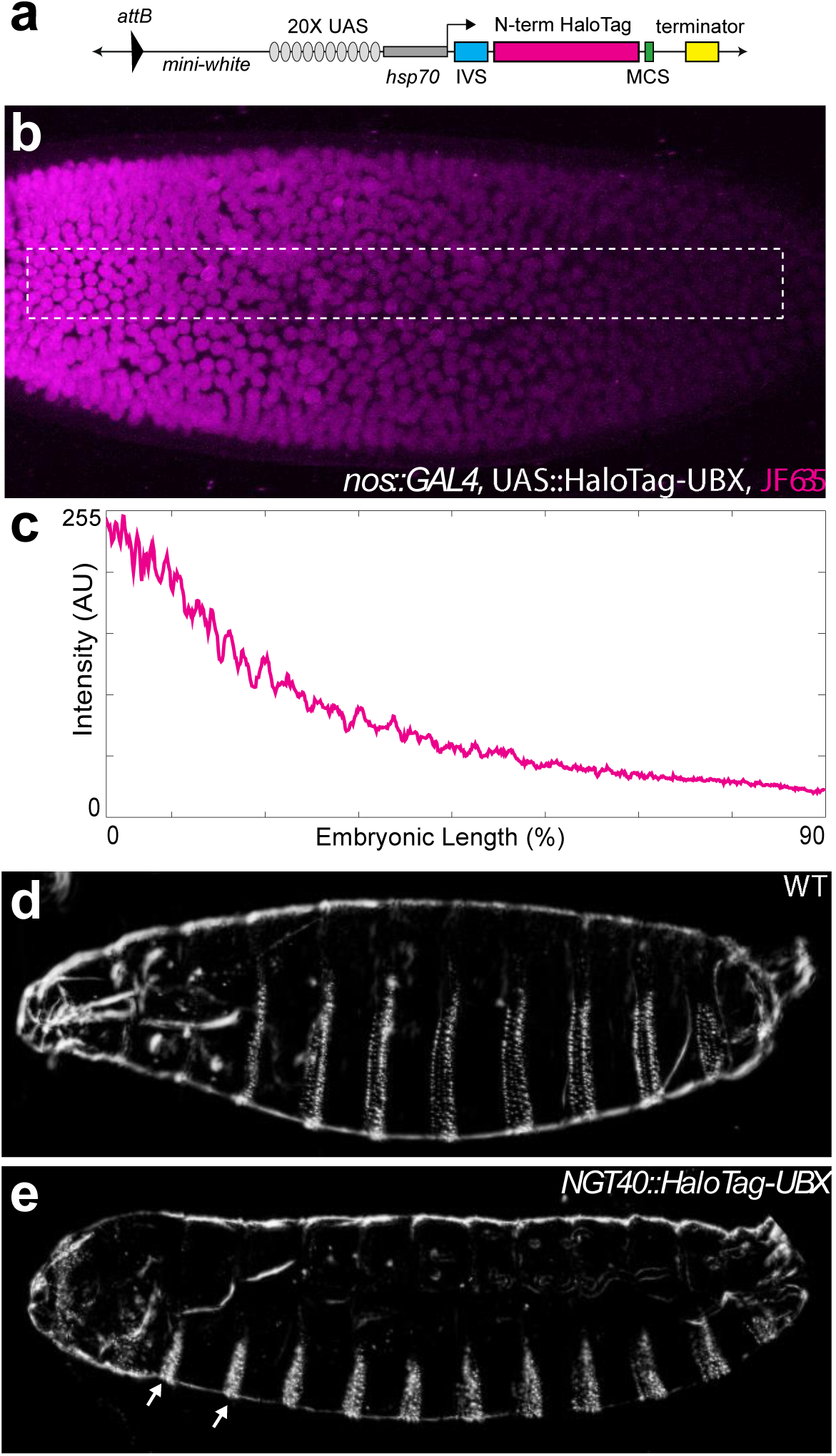
Control experiments for HaloTag-Ubx. (**a**) Schematic of the N-terminal HaloTag construct. (**b**) Stage 5 embryo with *nos*::*GAL4*, UAS::HaloTag-Ubx and injected with JF_635_ dye at the anterior end (magenta). (**c**) Quantification of the signal intensity of the bounded region in panel (**b**) along the embryonic axis. A.U. indicates Arbitrary Units of fluorescence intensity. (**d, e**) Cuticle preps of first instar larva from either a wild-type (WT) (**d**) or *nos*::GAL4, UAS::HaloTag-Ubx embryo. (**e**) Arrows point to the *nos*::GAL4, UAS::HaloTag-Ubx induced transformation of anterior segments.

**Figure S5:**
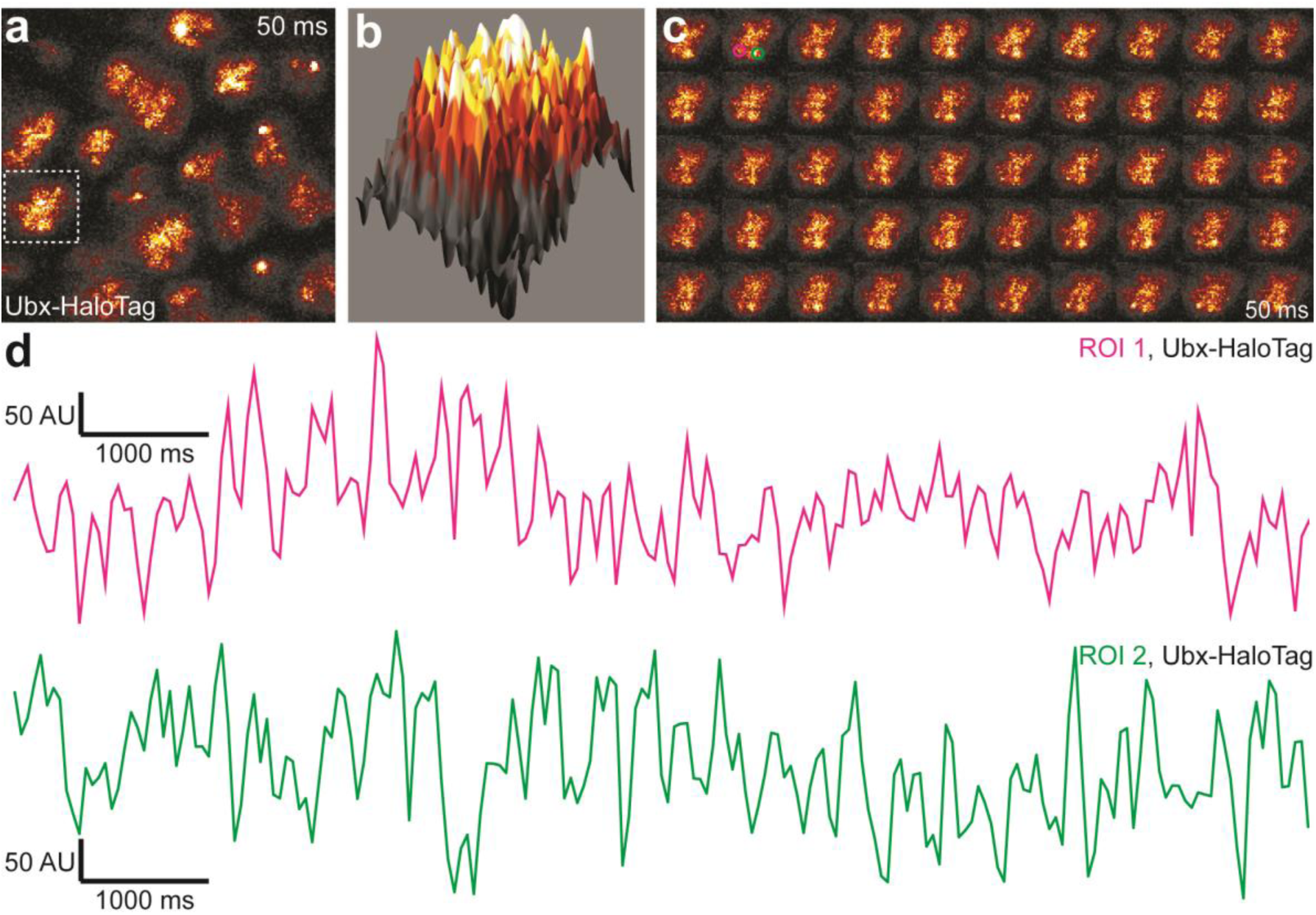
Live imaging multiple molecules of HaloTag-Ubx. (**a**) Fifty millisecond video frame from video of *nos*::GAL4, UAS::HaloTag-Ubx with a higher number of HaloTag-Ubx molecules (compare with Fig 1), imaged with JF_635_ dye. (**b**) 3D surface plot of the nucleus indicated in panel (**a**). (**c**) Individual, 50 millisecond video frames of the nucleus from panel (**a**). (**d**) Signal traces of the signal intensity of the regions noted with red and green circles in panel (**c**). AU indicates Arbitrary Units of fluorescence intensity.

**Figure S6:**
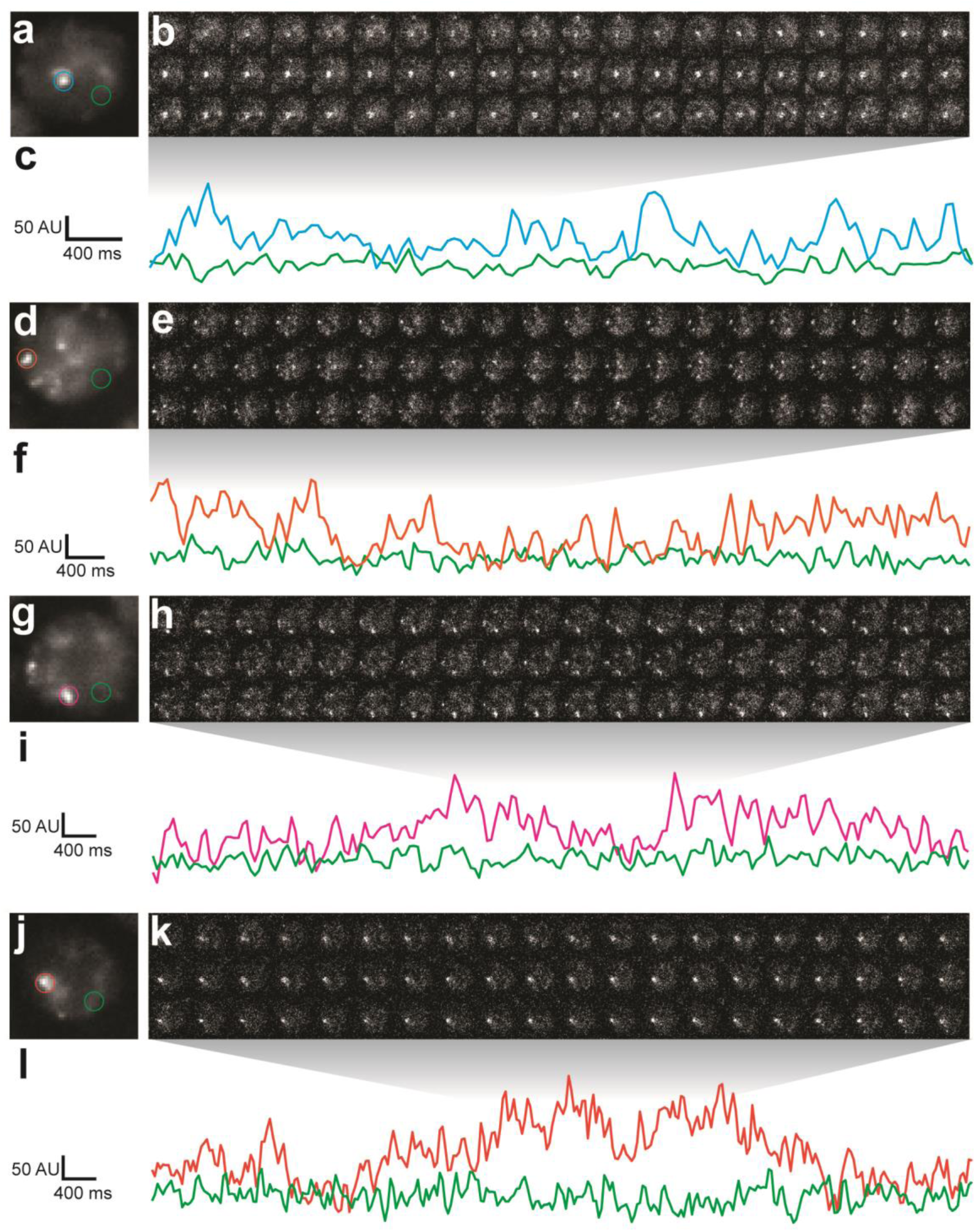
Additional nuclei from live imaging of Halo-Ubx. (**a, d, g, j**) Projections of summed pixel intensities over 100 seconds of *nos*::GAL4, UAS::HaloTag-Ubx nuclei imaged with JF_635_ dye. (**b, e, h, k**) Individual 50 millisecond video frames of the nuclei in panels (**a, d, g, j**). (**c, f, i, l**) Signal traces of the signal intensity of the regions noted in panels (**a, d, g, j**), where color of trace corresponds to color of circle. AU indicates Arbitrary Units of fluorescence intensity.

**Figure S7:**
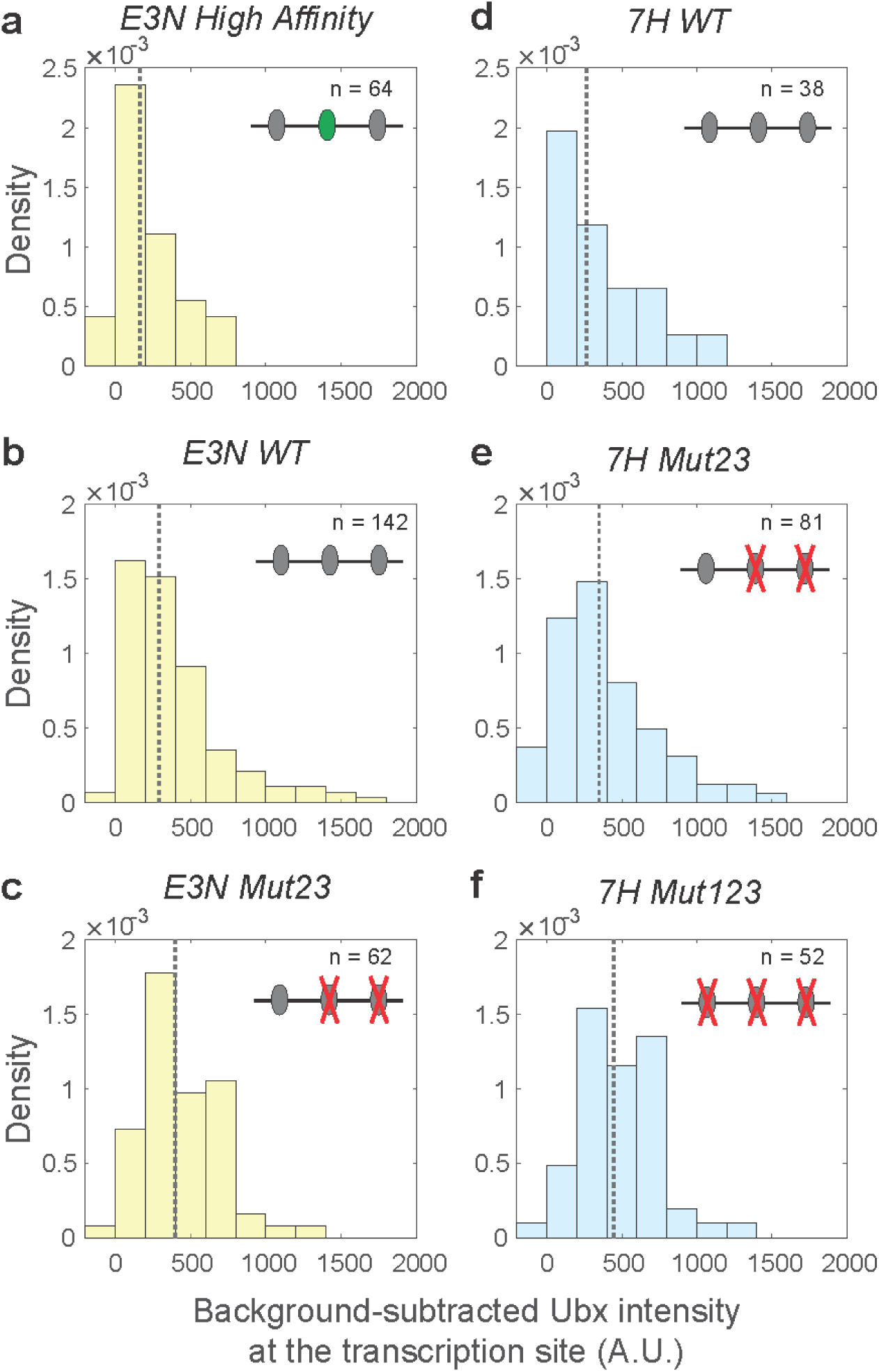
Background-subtracted Ubx intensity distributions at the transcription sites for *E3N* and *7H* enhancers. The Ubx intensity distributions at transcription sites were plotted for the *E3N* and *7H* enhancers after subtraction of background fluorescence from raw Ubx intensities. The naming convention of the enhancers follows that of Figure 3. The density (*y*- axis) for each distribution is calculated by the count per bin divided by a normalization factor. The normalization factor is the bin size multiplied by the number of transcription sites (*n*) in the dataset. The dotted gray line is the median of the distributions, which are: (**a**) 160 for *E3N High Affinity*, (**b**) 290 for *E3N WT*, (**c**) 400 for *E3N Mut23*, (**d**) 260 for *7H WT*, (**e**) 350 for *7H Mut23*, and (**f**) 440 for *7H Mut123*.

## References

1. Spitz, F. & Furlong, E. E. M. Transcription factors: from enhancer binding to developmental control. Nat. Rev. Genet. 13, 613–26 (2012).

2. Reiter, F., Wienerroither, S. & Stark, A. Combinatorial function of transcription factors and cofactors. Curr. Opin. Genet. Dev. 43, 73–81 (2017).

3. Long, H. K., Prescott, S. L. & Wysocka, J. Ever-Changing Landscapes: Transcriptional Enhancers in Development and Evolution. Cell 167, 1170–1187 (2016).

4. Crocker, J., Ilsley, G. R. & Stern, D. L. Quantitatively predictable control of Drosophila transcriptional enhancers in vivo with engineered transcription factors. Nat Genet 48, 292–298 (2016).

5. Liu, Z. et al. 3D imaging of Sox2 enhancer clusters in embryonic stem cells. Elife 3, e04236 (2014).

6. Chen, J. et al. Single-molecule dynamics of enhanceosome assembly in embryonic stem cells. Cell 156, 1274–1285 (2014).

7. Izeddin, I. et al. Single-molecule tracking in live cells reveals distinct target-search strategies of transcription factors in the nucleus. Elife 3, (2014).

8. Voss, T. C. et al. Dynamic Exchange at Regulatory Elements during Chromatin Remodeling Underlies Assisted Loading Mechanism. Cell 146, 544–554 (2011).

9. Normanno, D. et al. Probing the target search of DNA-binding proteins in mammalian cells using TetR as model searcher. Nat. Commun. 6, 7357 (2015).

10. Morisaki, T., Müller, W. G., Golob, N., Mazza, D. & McNally, J. G. Single-molecule analysis of transcription factor binding at transcription sites in live cells. Nat. Commun. 5, 4456 (2014).

11. Crocker, J. et al. Low affinity binding site clusters confer hox specificity and regulatory robustness. Cell 160, 191–203 (2015).

12. Farley, E. K. et al. Suboptimization of developmental enhancers. Science (80-.). 350, 325–328 (2015).

13. Farley, E. K., Olson, K. M., Zhang, W., Rokhsar, D. S. & Levine, M. S. Syntax compensates for poor binding sites to encode tissue specificity of developmental enhancers. Proc. Natl. Acad. Sci. 113, 6508–6513 (2016).

14. Lorberbaum, D. S. et al. An ancient yet flexible *cis*-regulatory architecture allows localized Hedgehog tuning by *patched/Ptch1*. Elife 5, (2016).

15. Antosova, B. et al. The Gene Regulatory Network of Lens Induction Is Wired through Meis-Dependent Shadow Enhancers of Pax6. PLOS Genet. 12, e1006441 (2016).

16. Rister, J. et al. Single-base pair differences in a shared motif determine differential Rhodopsin expression. Science 350, 1258–61 (2015).

17. Crocker, J., Potter, N. & Erives, A. Dynamic evolution of precise regulatory encodings creates the clustered site signature of enhancers. Nat. Commun. 1, 99 (2010).

18. Crocker, J., Noon, E. P. B. & Stern, D. L. The Soft Touch: Low-Affinity Transcription Factor Binding Sites in Development and Evolution. Curr. Top. Dev. Biol. (2015).

19. Tanay, A. Extensive low-affinity transcriptional interactions in the yeast genome. Genome Res. 16, 962–72 (2006).

20. Lebrecht, D. et al. Bicoid cooperative DNA binding is critical for embryonic patterning in Drosophila. Proc. Natl. Acad. Sci. U. S. A. 102, 13176–81 (2005).

21. Rowan, S. et al. Precise temporal control of the eye regulatory gene Pax6 via enhancer-binding site affinity. Genes Dev. 24, 980–5 (2010).

22. Gaudet, J. & Mango, S. E. Regulation of organogenesis by the Caenorhabditis elegans FoxA protein PHA-4. Science 295, 821–5 (2002).

23. Jiang, J. & Levine, M. Binding affinities and cooperative interactions with bHLH activators delimit threshold responses to the dorsal gradient morphogen. Cell 72, 741–752 (1993).

24. Ramos, A. I., Barolo, S. & B, P. T. R. S. Low-affinity transcription factor binding sites shape morphogen responses and enhancer evolution. Philos Trans R Soc L. B Biol Sci 368, 20130018 (2013).

25. Tillberg, P. W. et al. Protein-retention expansion microscopy of cells and tissues labeled using standard fluorescent proteins and antibodies. Nat Biotechnol 34, 987–992 (2016).

26. Teves, S. S. et al. A dynamic mode of mitotic bookmarking by transcription factors. Elife 5, (2016).

27. Grimm, J. B. et al. A general method to improve fluorophores for live-cell and single-molecule microscopy. Nat. Methods 12, 244–250 (2015).

28. Grimm, J. B. et al. A general method to fine-tune fluorophores for live-cell and in vivo imaging. bioRxiv (2017).

29. Gebhardt, J. C. M. et al. Single-molecule imaging of transcription factor binding to DNA in live mammalian cells. Nat. Methods 10, 421–426 (2013).

30. Slattery, M. et al. Cofactor binding evokes latent differences in DNA binding specificity between Hox proteins. Cell 147, 1270–82 (2011).

31. Crocker, J. & Stern, D. L. TALE-mediated modulation of transcriptional enhancers in vivo. Nat. Methods 10, 762–7 (2013).

32. Crocker, J., Tsai, A. & Stern, D. L. A Fully Synthetic Transcriptional Platform for a Multicellular Eukaryote. Cell Rep 18, 287–296 (2017).

33. Rieckhof, G. E., Casares, F., Ryoo, H. D., Abu-Shaar, M. & Mann, R. S. Nuclear translocation of extradenticle requires homothorax, which encodes an extradenticle-related homeodomain protein. Cell 91, 171–83 (1997).

34. Ryoo, H. D. & Mann, R. S. The control of trunk Hox specificity and activity by Extradenticle. Genes Dev. 13, 1704–16 (1999).

35. Lelli, K. M., Noro, B. & Mann, R. S. Variable motif utilization in homeotic selector (Hox)-cofactor complex formation controls specificity. Proc. Natl. Acad. Sci. U. S. A. 108, 21122–7 (2011).

36. Choo, S. W., White, R., Russell, S., Mi, H. & Diemer, K. Genome-Wide Analysis of the Binding of the Hox Protein Ultrabithorax and the Hox Cofactor Homothorax in Drosophila. PLoS One 6, e14778 (2011).

37. Slattery, M., Ma, L., Négre, N., White, K. P. & Mann, R. S. Genome-Wide Tissue-Specific Occupancy of the Hox Protein Ultrabithorax and Hox Cofactor Homothorax in Drosophila. PLoS One 6, e14686 (2011).

38. Dueber, J. E. et al. Synthetic protein scaffolds provide modular control over metabolic flux. Nat. Biotechnol. 27, 753–759 (2009).

39. Oehler, S. & Müller-Hill, B. High Local Concentration: A Fundamental Strategy of Life. J. Mol. Biol. 395, 242–253 (2010).

40. Yao, J., Munson, K. M., Webb, W. W. & Lis, J. T. Dynamics of heat shock factor association with native gene loci in living cells. Nature 442, 1050–1053 (2006).

41. Zhang, C. et al. A clustering property of highly-degenerate transcription factor binding sites in the mammalian genome. Nucleic Acids Res. 34, 2238–46 (2006).

42. Elf, J., Li, G.-W. & Xie, X. S. Probing transcription factor dynamics at the single-molecule level in a living cell. Science 316, 1191–4 (2007).

43. Kabata, H. et al. Visualization of single molecules of RNA polymerase sliding along DNA. Science (80-.). 262, 1561–1563 (1993).

44. Leith, J. S. et al. Sequence-dependent sliding kinetics of p53. Proc. Natl. Acad. Sci. U. S. A. 109, 16552–7 (2012).

45. Ruusala, T. & Crothers, D. M. Sliding and intermolecular transfer of the lac repressor: kinetic perturbation of a reaction intermediate by a distant DNA sequence. Proc. Natl. Acad. Sci. U. S. A. 89, 4903–7 (1992).

46. Junion, G. et al. A transcription factor collective defines cardiac cell fate and reflects lineage history. Cell 148, 473–86 (2012).

47. Noordermeer, D. et al. Temporal dynamics and developmental memory of 3D chromatin architecture at Hox gene loci. Elife 3, e02557 (2014).

48. de Laat, W. & Duboule, D. Topology of mammalian developmental enhancers and their regulatory landscapes. Nature 502, 499–506 (2013).

49. Symmons, O. et al. The Shh Topological Domain Facilitates the Action of Remote Enhancers by Reducing the Effects of Genomic Distances. Dev. Cell 39, 529–543 (2016).

50. Williamson, I., Lettice, L. A., Hill, R. E. & Bickmore, W. A. Shh and ZRS enhancer colocalisation is specific to the zone of polarising activity. Development 143, 2994–3001 (2016).

51. Giorgetti, L. et al. Structural organization of the inactive X chromosome in the mouse. Nature 535, 575–579 (2016).

52. Fukaya, T., Lim, B. & Levine, M. Enhancer Control of Transcriptional Bursting. Cell 166, 358–368 (2016).

53. Cisse, I. I. et al. Real-Time Dynamics of RNA Polymerase II Clustering in Live Human Cells. Science (80-.). 341, 664–667 (2013).

54. Cho, W.-K. et al. RNA Polymerase II cluster dynamics predict mRNA output in living cells. Elife 5, 681–686 (2016).

55. Boettiger, A. N. et al. Super-resolution imaging reveals distinct chromatin folding for different epigenetic states. Nature 529, 418–22 (2016).

56. Hnisz, D., Shrinivas, K., Young, R. A., Chakraborty, A. K. & Sharp, P. A. A Phase Separation Model for Transcriptional Control. Cell 169, 13–23 (2017).

57. Sheppard, C. J., Mehta, S. B. & Heintzmann, R. Superresolution by image scanning microscopy using pixel reassignment. Opt Lett 38, 2889–2892 (2013).

58. Schindelin, J., Rueden, C. T., Hiner, M. C. & Eliceiri, K. W. The ImageJ ecosystem: An open platform for biomedical image analysis. Mol Reprod Dev 82, 518–529 (2015).

59. Groth, A. C., Fish, M., Nusse, R. & Calos, M. P. Construction of transgenic Drosophila by using the site-specific integrase from phage phiC31. Genetics 166, 1775–1782

60. Schindelin, J. et al. Fiji: an open-source platform for biological-image analysis. Nat Methods 9, 676–682 (2012).

61. Schmid, B., Schindelin, J., Cardona, A., Longair, M. & Heisenberg, M. A high-level 3D visualization API for Java and ImageJ. BMC Bioinformatics 11, 274 (2010).

